# Rab18 Binds PLIN2 and ACSL3 to Mediate Lipid Droplet Dynamics

**DOI:** 10.1101/2020.05.02.073957

**Authors:** Yaqin Deng, Chang Zhou, Mirza Ahmed Hammad, Adekunle T. Bamigbade, Shimeng Xu, Shuyan Zhang, Pingsheng Liu

**Affiliations:** National Laboratory of Biomacromolecules, CAS Center for Excellence in Biomacromolecules, Institute of Biophysics, Chinese Academy of Sciences, Beijing, 100101, China; University of Chinese Academy of Sciences, Beijing, 100049, China

**Keywords:** Lipid droplets, Rab18, PLIN2, ACSL3, TAG

## Abstract

Rab18 has been linked to lipid metabolism and metabolic diseases in different model systems, but the mechanism of Rab18-mediated lipid droplet (LD) dynamics in muscle cells remains elusive. Here, we report that Rab18 plays an essential role in oleic acid (OA)-induced LD growth and formation in mouse myoblast cell line C2C12. Rab18 was translocated from endoplasmic reticulum (ER) to LDs during the LD growth induced by OA in C2C12 cells, which was directly regulated by perilipin 2 (PLIN2), a LD resident protein. LD-associated Rab18 bound with the C terminus of PLIN2, and the LD localization of Rab18 was diminished after PLIN2 deletion. Moreover, loss of function of Rab18 led to less triacylglycerol (TAG) accumulation and fewer but larger LD formation. In contrast, expression of wild type Rab18 and a constitutively active Rab18 (Q67L) mutant resulted in elevated TAG content and LD number. Furthermore, LD-associated Rab18 interacted with acyl-CoA synthetase long-chain family member 3 (ACSL3) and in turn, promoted the LD localization of ACSL3, which may play an important role in the accumulation of TAG induced by OA. These data showed that Rab18 was recruited to LD after OA treatment, and formed a complex with PLIN2 and ACSL3, which contributes to TAG accumulating and LD growth.

## Introduction

Lipid droplets (LDs) are a dynamic organelle that widely exists in bacteria, yeast, plants, *C. elegans*, insects, and mammalian cells [1, 2]. It consists of neutral lipid core and phospholipid monolayer membrane which is coated by various types of proteins [3–5]. In addition to serving as energy storage reservoir, LDs are found to be involved in several processes, including lipid synthesis and metabolism, membrane trafficking, protein storage and degradation, and cell signaling [6, 7]. The number and size of LDs are closely related to many metabolic diseases, such as obesity, type 2 diabetes, fatty liver, and lipodystrophies [8, 9].

The diversity of LD functions greatly depends on the proteins associated with the organelle. The perilipin family proteins (PLINs) play important roles in LD biogenesis and functions in mammalian cells [10]. PLIN1, PLIN2 and PLIN5 are involved in lipolysis which is coordinated with the recruitment of cytoplasmic lipases [11–14]. Other lipid metabolism-related enzymes are also found to be localized on LDs, which can promote LD biogenesis or triacylglycerol (TAG) accumulation and hydrolysis, including acyl-CoA synthetase long chain family member 3 (ACSL3), diglyceride acyltransferase1/2 (DGAT1/2), hormone sensitive lipase (HSL), and adipose triglyceride lipase (ATGL) [15–18]. Besides, nearly 30 Rab GTPases are identified in different proteomes of isolated LDs [19–22]. However, just a small part of these Rab proteins have been proved to localize on LDs through at least two different methods or closely be related to the function of LDs, including Rab1, Rab5, Rab7, Rab8a, Rab18, Rab32 and Rab40c [23]. Rab proteins are the major regulators of membrane trafficking and circulate between active form (GTP-bound) and inactive form (GDP-bound). The LD-associated Rab proteins mainly take part in LD formation, LD fusion and the interactions between LD and other organelles [24–27].

Rab18 is involved in many biological processes including secretion inhibiting, Golgi and endoplasmic reticulum (ER) membrane trafficking, ER morphology maintaining, macroautophagy promoting, LD dynamic regulating, and lipid metabolism [26, 28–35]. The co-localization of Rab18 and LDs was fully proved by fluorescence imaging, immunogold electron microscope imaging, and subcellular fractionation [26, 27, 33, 34]. Rab18 has been shown to mediate the interaction between LD and ER [26, 33]. Li et al. also reveals that the DFCP1-Rab18 complex is involved in tethering the ER-LD contact for LD expansion and initial LD growth [36, 37]. Another possible mechanism is that Rab18 specifically binds to the ER-associated NAG-RINT1-ZW10 tethering complex [38]. In adipocytes, Rab18 may play an important role in both TAG synthesis and hydrolysis [33, 34]. It has been reported that the LD targeting of Rab18 is regulated by its GEF TRAPPII and Rab3GAP1/2 in Huh7 cells and 3T3-L1 pre-adipocytes [38, 39]. However, even in TRAPPII or Rab3GAP1/2 deficient cells, there is still some LD-associated Rab18. It suggests that the LD association of Rab18 is controlled by multiple mechanisms. Loss-of-function mutations in Rab18 itself, its GEF protein Rab3GAP1/2 and its GAP protein TBC1D20 cause Warburg Micro syndrome (WARBM), which implies the direct molecular mechanism in the disease pathology [40–44]. Interestingly, in all WARBM fibroblasts with mutations in these three genes and in MEFs from Rab18 or TBC1D20 deletion mouse models, aberrant LD formation is a cellular common phenotype [43, 45]. Aberrant lipid storage in LDs is the key factor of ectopic lipid accumulation that has been linked to the progression of many metabolic disorders. Initially the LD targeting phenotype of Rab18 leads us to the relationship between Rab18 and LD dynamics. Recently, Xu et al. [38] claimed that Rab18 controls the growth of LD, but does not affect the LD biogenesis by tracking the formation of nascent LD in Rab18 deficient cells. However, more evidence needs to elucidate the specific mechanism.

In this study, we investigated the potential role of Rab18 in LD dynamics in myoblasts. We found that with the LD growth and TAG accumulation, more Rab18 was recruited to LDs. The translocation of Rab18 from ER to LDs was mediated by the LD resident protein, PLIN2, in C2C12 cells. The LD-associated Rab18 directly interacted with the C-terminal of PLIN2. At the same time, Rab18 recruited ACSL3 to LDs, formed a PLIN2/Rab18/ACSL3 complex, that in turn facilitated LDs activity, which contributes to TAG accumulation and LD growth.

## Materials and methods

### Reagents, plasmids and antibodies

Hoechst 33342, LipidTOX Red, Lipofectamine 2000, OptiPrep and Puromycin were purchased from Invitrogen. Anti-FLAG-M2 beads, DMSO, Sodium oleate, PMSF, Polybrene and Triton X-100 were from Sigma. DSP was from Thermo.

cDNAs encoding Rab18, PLIN2, ACSL3 and Sec61β were amplified by PCR from cDNAs of C2C12 cells. Rab18 was cloned into pEGFP-C1. FLAG-tagged Rab18 was cloned into pQCXIP. Rab18-S22N and Rab18-Q67L mutations were constructed by a PCR-based site-directed mutagenesis. Sec61β was cloned into pmCherry-C1, which was modified from pEGFP-C1 by displacing GFP with mCherry. PLIN2 was cloned into pFLAG-CMV4. PLIN2 was cloned into pEGFP-N1, as well as its truncations. FLAG-tagged ACSL3 was cloned into pQCXIP.

Anti-Rab18 (Rabbit, A2812), anti-Tip47 (Rabbit, A6822) and anti-PDI (Rabbit, A0692) antibodies were purchased from ABclonal. Anti-PLIN2 antibody (Rabbit, ab108323) was from Abcam. Anti-LAMP1 antibody (Mouse, 3243s) was from Cell Signaling Technology. Anti-Bip (Mouse, 610979), anti-GM130 (Mouse, 610822) and anti-Tim23 (Mouse, 611223) antibodies were from BD Transduction. Anti-GAPDH antibody (Mouse, MAB374) was from Millipore. Anti-Actin antibody was from Huaxinbio (Mouse, HX1827). Anti-GFP antibody was from Santa Cruz (Mouse, sc-9996). Anti-ACSL3 antibody was from Proteintech (Rabbit, 20710-1-AP).

### Cell culture, transfection and RNAi interference

Mouse C2C12 myoblasts (American Type Culture Collections), HEK293A and Plat-ET cell lines were maintained in DMEM (Macgene Biotechnology, CN) supplemented with 10% FBS (Gibco) and 100 U/mL penicillin and 100 mg/mL streptomycin

(Macgene Biotechnology, CN) at 37°C in a humidified 5% CO_2_ incubator. C2C12 cells was treated with or without 100 μM OA for 12 h. C2C12 cells were transfected with indicated plasmids using electroporation according to the manufacturer’s instructions (Amaxa Nucleofector). For siRNA transfection, unless otherwise indicated, all the experiments were performed at 72 h after the transfection. The sequences of each siRNA oligonucleotides (GenePharm, CN) target to mouse Rab18 are as follows: Rab18-siRNA-1: 5’-GGCUAAACUUGCAAUAUGG-3’; Rab18-siRNA-2: 5’-ACCGUGAAGUCGAUAGAAA-3’. HEK293A and Plat-ET cells were transfected by using Lipofectamine2000 according to the manufacturer’s instructions.

### CRISPR/Cas9 knockout

Two targets corresponding separately to Exon 1 and Exon 4 of Rab18 were designed according to website (http://crispr.mit.edu/) and constructed into pX260a plasmid (gifted from Prof. Feng Zhang [46]). The sequences of the two guide RNA are as follows: Rab18-KO-1: 5’-TACTCATCATCGGCGAGAGT-3’; Rab18-KO-2: 5’-CTGTGCACCTCTATAATAGC-3’. The two plasmids were transfected into C2C12 cells in a 6-well plate. After 48 h, the transfected cells were expanded into a 10-cm plate and selected with growth medium containing 1 μg/mL puromycin. After 2 weeks, the puromycin-resistant cells were then seeded into 96-well plates with gradient dilution and cultured for another 2 weeks before picking the monoclones. The picked monoclones were verified by Western blots.

Plin2 KO cells were generated in our lab as described previously [47].

### Stable overexpression cell line construction

The coding sequence of indicated genes was cloned into a retroviral expression vector, pQCXIP. The pQCXIP derived plasmids, combined with two virus structural genes VSV-G and Phit, were packaged into pseudo retrovirus within Plat-ET cells. 48 h after transfection, the viruses were collected and used to infect C2C12 cells in a 6-well plate with 10 μg/mL polybrene. Then the infected cells were expanded into a 10-cm plate and selected with growth medium containing 1 μg/mL puromycin. The puromycin-resistant cells were screened for monoclones.

### LD isolation

LDs were isolated by a modified method of Zhang et al[48]. Briefly, C2C12 cells with the indicated treatments were rinsed with ice-cold PBS and then pelleted. The pellets were suspended in 10 mL Buffer A (25 mM Tricine, pH 7.8, 250 mM Sucrose) containing 0.5 mM PMSF and incubated on ice for 20 min. The swelling cells were then homogenized by N_2_ bomb (500 psi for 15 min on ice). Cell debris was removed by centrifugation at 3,000*g* for 10 min at 4°C. The post-nuclear supernatant (PNS) fraction (8 mL) was collected and loaded into a SW40 tube which was further overlaid with about 3 mL Buffer B (20 mM HEPES, pH 7.4, 100 mM KCl, and 2 mM MgCl_2_). The sample was centrifuged at 250,000*g* for 1 h at 4°C. The white band containing LDs at the top of gradient was collected into a 1.5 mL Eppendorf tube and washed three times with Buffer B by centrifuging at 20,000*g* at 4°C. Finally, the LD proteins were precipitated with 1 mL acetone treatment followed by centrifuging at 20,000*g* for 10 min at 4°C. The protein pellet was then dissolved in 2 × sample buffer (125 mM Tris Base, 20% Glycerol, 4% SDS, 4% β-mercaptoethanol and 0.04% Bromophenol blue). Following the ultracentrifugation step, the pellet was collected as total membrane (TM) and the medium layer was collected as cytosol (Cyto).

### Cell fractionation

The indicated cells were rinsed with ice-cold PBS and then pelleted. The pellets were suspended in 3 mL Buffer A (25 mM Tricine, pH 7.8, 250 mM Sucrose) containing with 0.5 mM PMSF and then incubated on ice for 20 min. The swelling cells were then homogenized by N_2_ bomb (500 psi for 15 min on ice). Cell debris was removed by centrifugation at 3,000*g* for 10 min at 4°C. The supernatant was collected, and the protein concentration was measured by BCA. 5 mg proteins in 2 mL was mixed with 2 mL 60% OptiPrep, which was loaded in the SW40 tubes. 4 mL 20% OptiPrep and 5 mL 10% OptiPrep were loaded into the samples separately and slowly. The sample was centrifuged at 35,000 rpm for 18 h at 4°C. Take 1 mL liquid from the top to bottom each time as different fractions. All the fractions were performed TCA precipitation, and then dissolved in 2 × sample buffer.

### TAG measurement

The indicated cells were inoculated into 6 well plates. After OA treatment, the cells were lysed in 1% Triton X-100. The cell extracts were used to determine the TAG content by TAG kit (GPO-PAP method, BioSino Bio) and total protein content by BCA assay (Thermo).

### Western blot analysis

The indicated proteins were loaded to SDS-PAGE gel and transferred to methanol-activated PVDF membranes (Millipore), which were blocked in 5% non-fat milk containing 0.2% Tween for 1 h at room temperature (RT). Then, the membranes were incubated with primary antibodies for 1 h at RT or overnight at 4°C. After 1 h incubation with secondary antibodies at RT, the membranes were exposed with the ECL substrate (PerkinElmer Life Sciences).

### Immunoprecipitation

For LD-IP, LDs were isolated as describe above, and lysed in 0.5 mL immunoprecipitation (IP) buffer (50 mM Tris-HCl, pH 7.5, 150 mM NaCl, 1 mM EDTA, 1% Triton X-100, 10% Glycerol) with supplied protease inhibitor cocktail (CWBiotech) for 30 min at 4°C. Anti-FLAG M2 beads were blocked with IP buffer supplied 200 μg BSA for three times. The sample was centrifuged at 20,000*g* for 5 min at 4°C. The clear liquid below the LDs was collected and incubated with blocked beads at 4°C for 3 h. The beads were washed with IP buffer for three times, and then the proteins on the beads were dissolved in 2 × sample buffer.

For whole cell lysate IP, after transfection or OA treatment, cross linking was conducted with DSP according to the manufacturer’s instructions. Then the cells were lysed and proteins IP as above.

### Immunofluorescence

The cells were cultured on coverslips and treated with OA. After washed with PBS for three times, the cells were fixed with 4% paraformaldehyde for 30 min and permeabilized with 0.01% Digitonin for 30 min. Then the cells were blocked with 1% BSA for 1 h, and incubated with anti-ACSL3 for 1 h, followed by incubation with Alexa Fluor 488 conjugated anti-rabbit secondary antibody (Thermo) for 1 h, and then stained with LipidTOX Red and Hoechst. After mounted, pictures were captured by laser scanning confocal microscope.

### Bioinformatic analysis

Rab18 structure was generated from I-tasser web server with PDB ID 1X3S, while ADRP/PLIN2 model was generated through I-tasser web server without providing template structure. Both models were minimized prior to be used in Protein-Protein docking experiment. Minimization removes energy constraints from protein 3D models making them more flexible. For minimization, UCSF Chimera tool was utilized with 1000 steepest descent and 1000 conjugate gradient steps. The minimized proteins were subjected to PDBsum tool available on www.ebi.ac.uk. PDBsum calculates different protein features including secondary structure elements and Ramachandran values. The proteins were then subjected to protein-protein docking through two web servers; i) PatchDock, ii) GrammX. The results from both tools were then refined by FireDock tool which refined protein-protein docking solutions and also calculated their global energy for interacting residues.

### Statistical analyses

Data were presented as mean ± SD. The statistical analyses were performed using Graphpad Prism 6. Comparison of significance was assessed by two-tailed Student’s *t* test. For all analyses, a p-value < 0.05 was considered to be statistically significant.

## Results

### Recruitment of Rab18 from ER to LDs during OA-induced LD growth

To investigate LD dynamics and lipid homeostasis in myoblast, oleic acid (OA) was used to induce LD formation in C2C12 cells. The LD growing was gradually observed with the OA loading (Fig. 1A, 0-12 h). Rab18, a membrane trafficking GTPase, appeared and was significantly accumulated on the growing LDs (Fig. 1A). To examine how and why Rab18 was accumulated on OA-induced growing LDs in myoblast, the expression of Rab18 in whole cell lysate was firstly tested. Compared to significant increase of PLIN2, no change was detected after OA treatment for different times (Fig. 1B, lanes 1-4), suggesting that the localization rather than the expression level of Rab18 was altered by OA treatment. Since Rab18 was previously identified to be localized between ER and LDs by immunogold-EM [18, 24], an ER marker protein Sec61β was co-transfected with Rab18 to determine if the LD-associated Rab18 was translocated from ER. Result in Figure 1C showed that Rab18 was co-localized with Sec61β strongly before OA treatment, while the co-localization of these two proteins was disrupted by OA treatment.

**Figure 1.**
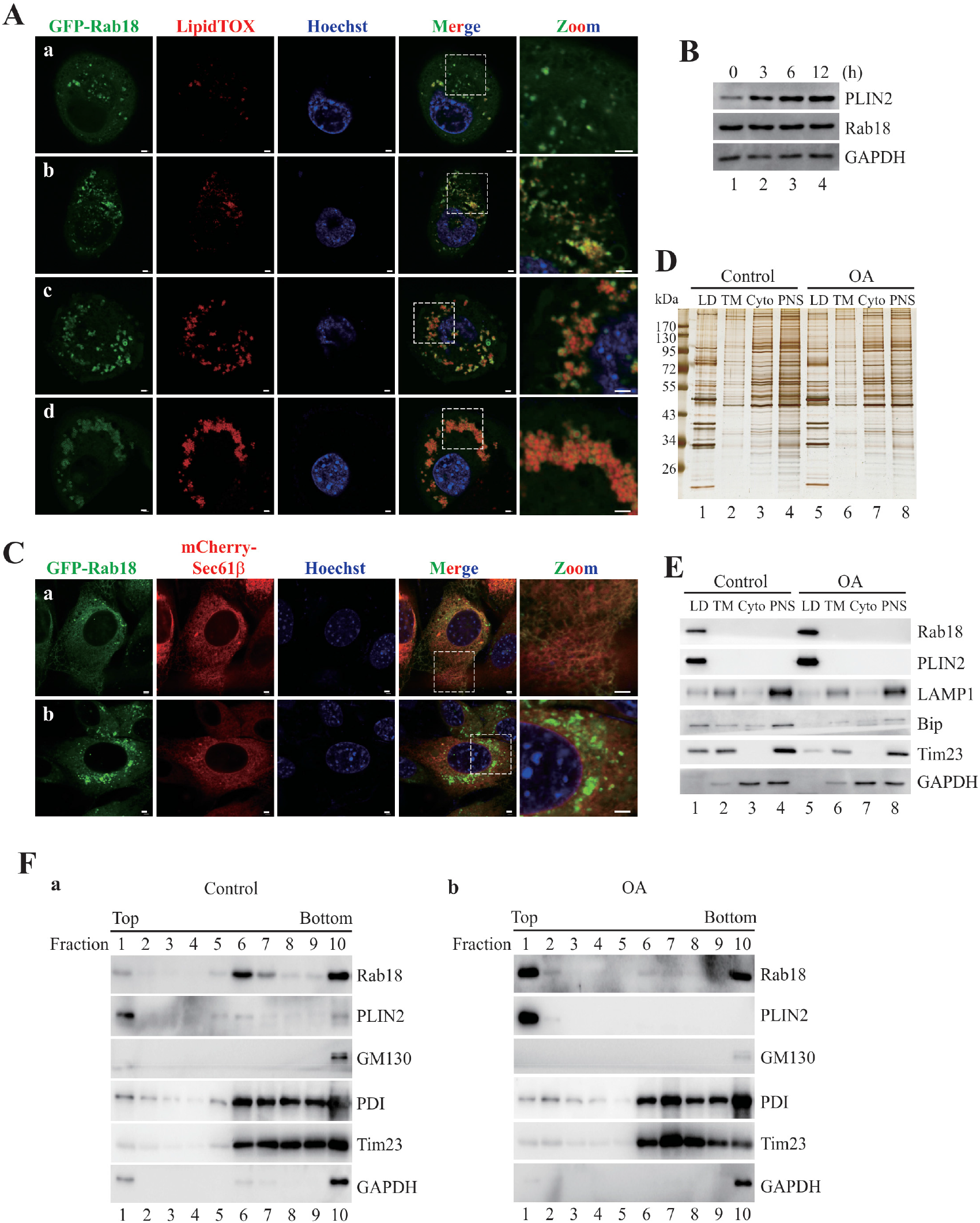
Recruitment of Rab18 from ER to LDs by OA treatment. **A**. C2C12 cells transiently transfected with GFP-Rab18, was treated with 100 μM OA, and colocalization of GFP-Rab18 and LD (lipidTOX) was acquired at 0, 3, 6, 12 hrs (panel **a-d**). Scale bar, 2 μm. **B**. Proteins expression level in C2C12 cells after OA treatment for indicated time. C. C2C12 cells were co-transfected with GFP-Rab18 and mCherry-Sec6β1. The colocalization of Rab18 and Sec61β before (**a**) and after (**b**) OA treatment, was imaged by using confocal microscope. Scale bar, 2 μm. **D-E**. LDs were isolated from C2C12 cells with or without 12 h OA treatment, then sliver staining (**D**) and Western bolt (**E**) were performed for the LD proteins, total membrane (TM) proteins, cytosolic (Cyto) proteins and post-nuclear supernatant (PNS). **F**. C2C12 cells without (**a**) or with (**b**) 12 h OA treatment, were fractionated by Opti-prep discontinuous density gradient centrifugation. The presence of Rab18 in the fractions was detected by immunoblotting, as well as LD marker PLIN2, Golgi marker GM130, ER marker PDI, mitochondrion marker Tim23 and cytosol marker GAPDH.

To explore the localization of endogenous Rab18, we isolated LDs from C2C12 cells with or without OA treatment (Fig. 1D). The purity of LDs was assessed by the distribution of different organelle marker proteins, including LD resident protein PLIN2, lysosome protein LAMP1, ER protein Bip, mitochondrion protein Tim23, and cytosol protein GAPDH. With similar protein amount loading, silver staining presented a unique protein composition of isolated LDs (LD) for both conditions, compared with total membrane (TM), cytosol (Cyto), and post nuclear supernatant (PNS) (Fig. 1D). Western blots of LDs and other cellular fractions showed that Rab18 in LDs was increased by OA treatment (Fig. 1E, lanes 1 and 5), based on equal protein loading (Fig. 1D, lanes 1 and 5). To investigate the translocation of Rab18 between two compartments in C2C12 cell with OA treatment, cellular organelles were fractionated by an Opti-prep discontinuous gradient, in which different organelles were distributed in different density fractions (Fig. 1F). In control cells (Control) as panel a shown, Rab18 were mainly detected in the fractions 6 and 10 that were enriched with ER protein PDI and mitochondrion protein Tim23. However, after OA treatment (OA), Rab18 was significantly transferred to the top fraction that merely contained the LD resident protein PLIN2 without increasing of ER protein PDI (panel b, lane 1). These results demonstrated that in myoblast cells Rab18 was translocated from ER to LDs during OA-induced LD growth.

### PLIN2 enhances Rab18 translocation from ER to LDs

Rab18 was identified to be associated with LDs by isolation plus proteomics in 2004 [26, 27, 33, 34] and then proved to be localized to LDs using immunogold-EM in 2007 [27]. Its function has been studied in different tissues and cell lines since then [38], however, the mechanism how Rab18 was recruited to LDs still need to be illuminated. Membrane targeting of Rab proteins is regulated by several factors including insertion of the modified C-terminus into the membrane and interaction with membrane proteins. Liu’s study suggested that Rab18 localized to LD by a different mechanism rather than simply C-terminus post-translational modification [27]. In 3T3-L1 preadipocytes, Rab3GAP1/2 and TRAPPII functioned as GEFs of Rab18, are required for LD association of Rab18 [38, 39].

To investigate the mechanisms of Rab18 translocation and the contributions to LD growth in myoblasts, we focused on its interacting proteins on LDs. LDs were isolated from FLAG-Rab18 overexpressed (OE) cells after OA treatment, and then the LD proteins were proceeded for immunoprecipitation (IP) by anti-FLAG M2 beads. The precipitates were then separated by SDS-PAGE and analyzed using sliver staining (Fig. 2A). Compared to control, three main distinct bands in FLAG-Rab18 IP samples were excised for mass spectrometry analysis. Among the proteins detected listed in Table 1, PLIN2 was found as the most abundant protein in both band 2 and 3, which is the main LD protein of C2C12 cells. Western blotting demonstrated that endogenous PLIN2 was significantly pulled-down in the precipitates, as well as ACSL3 (Fig. 2B). To verify this finding, IP was facilitated in a reverse way on FLAG-PLIN2 overexpressed C2C12 cells. The endogenous Rab18 could also be co-precipitated by FLAG-PLIN2 from OA-induced LD proteins (Fig. 2C). These results indicated that Rab18 and PLIN2 strongly interacted with each other on LDs. Since PLIN2 is a resident protein on LD, to identify whether PLIN2 contributes to the LD localization of Rab18, GFP-Rab18 was transiently transfected into Plin2 KO C2C12 cells. In WT control cells, almost all GFP-Rab18 was co-localized with LDs after OA treatment. However, when Plin2 was deleted, the LD localization of Rab18 was totally destroyed, and the GFP-Rab18 distributed pattern was similar in untreated WT cells (Fig. 2D). Furtherly, LDs from Plin2 KO cells were isolated and analyzed by Western blot and silver staining. As it shown in Fig. 2E and Fig. S1B, LD associated Rab18 was strikingly decreased when Plin2 was deleted, while the protein expression level of Rab18 was unchanged (Fig. 2F). Furthermore, subcellular organelles from WT and Plin2 KO cells were fractionated by density gradient centrifugation after treated with OA (Fig. 2G). In WT cells as panel a shown, endogenous Rab18 was mainly in the top two fraction which indicated to LD fractions, however, in Plin2 KO cells (panel b), the majority of Rab18 was in the fraction 6, which mainly contained ER marker protein PDI. Overall, these data strongly indicate that the interaction between Rab18 and PLIN2 is essential for the stabilization of Rab18 attachment to LDs.

**Figure 2.**
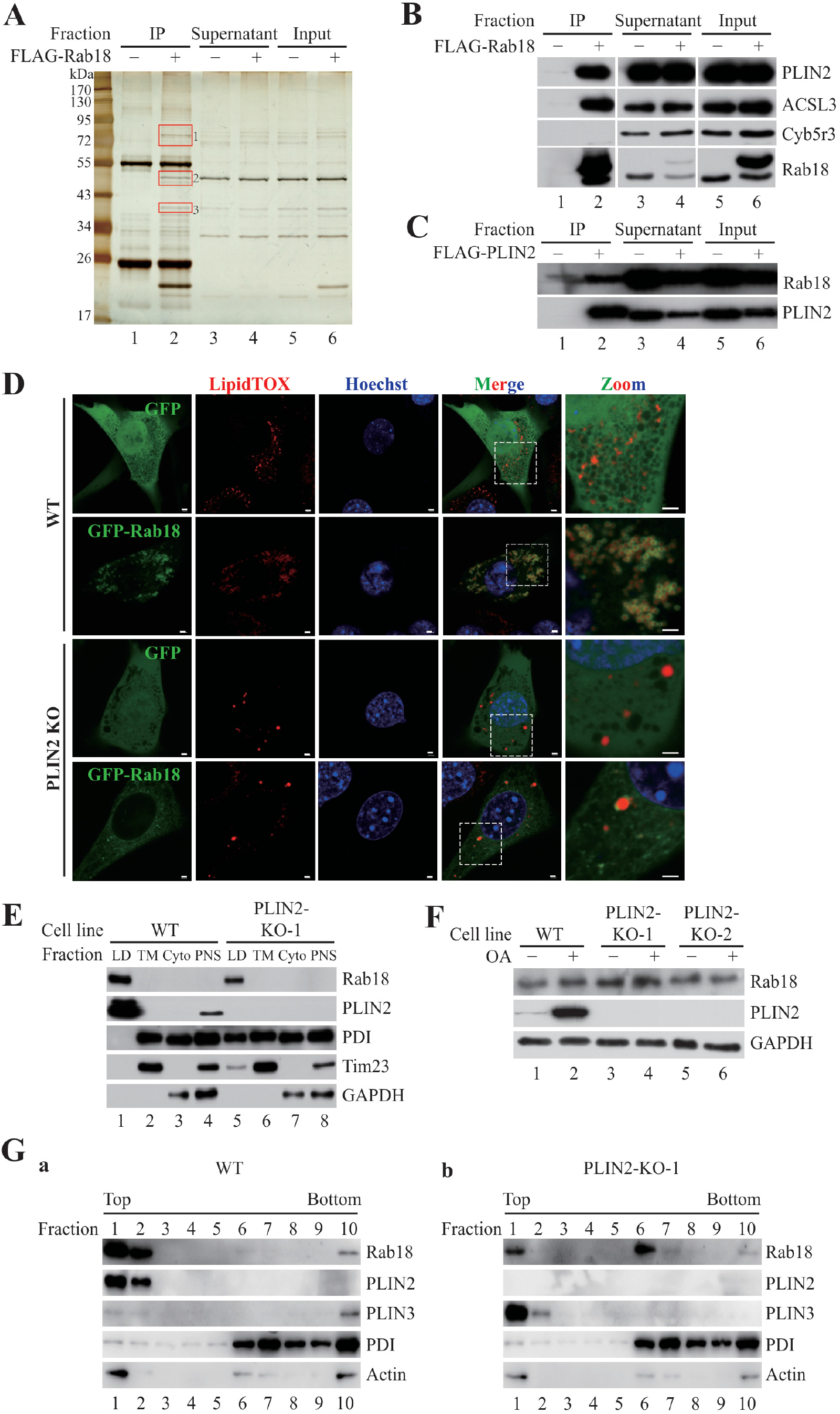
Rab18 translocation from ER to LD through interacting with PLIN2. **A**. LDs were isolated from OA treated C2C12 cells, including WT cells and FLAG-Rab18 overexpressed cells. The LD proteins were performed IP with anti-FLAG M2 beads then analyzed by silver staining. The three bands, specifically bound to FLAG-Rab18, were excised for mass spectrometry analysis. **B**. Western blotting analysis of the precipitates pulled-down by FLAG-Rab18. **C**. LD isolation was proceeded on OA treated C2C12 cells overexpressed with FLAG-PLIN2. LD proteins were lysed for IP with anti-FLAG M2 beads, then the precipitates were analyzed by Western blot. **D**. WT and Plin2-KO C2C12 cells were transiently transfected with GFP vector or GFP-Rab18, after treated with OA for 12h, the co-localization of Rab18 and LDs was imaged by confocal. Scale bar, 2 μm. **E**. LDs were isolated from C2C12 WT and Plin2-KO cells. With equal amount of protein loaded, LD associated Rab18 was detected by Western blot. **F**. Intracellular Rab18 Expression in two Plin2-KO monoclonal cells. **G**. After 12 h OA treatment, WT (**a**) and Plin2-KO-1 (**b**) C2C12 cells were fractionated by OptiPrep discontinuous density gradient centrifugation. The intracellular distribution of Rab18 was detected by Western blot. Fraction samples were equal volume loaded.

### The C terminal LD targeting domain of PLIN2 is required for interaction with Rab18

To investigate whether a direct interacting exists between PLIN2 and Rab18, the two-protein docking was predicted and analyzed by bioinformatics methods firstly (Fig. 3A). The docking results indicated PLIN2-Rab18 interacting residues mainly existed in the C terminus of PLIN2 (Table 2). According to the reported domains and predicted secondary structure of PLIN2 [49], PLIN2 was truncated into several parts (Fig. 3B). Co-IP between Rab18 and PLIN2 truncations was performed in HEK293A cells. Significant amounts of full length GFP-PLIN2 (GFP-PLIN2-FL) and GFP-PLIN2-251-425 were co-precipitated with FLAG-Rab18, while GFP-PLIN2-1-251 was not (Fig. 3C). Then PLIN2-251-425 was furtherly truncated into small parts on the basis of the interacting residues distribution (Fig. 3D). Only PLIN2-395-425 and PLIN2-370-425 could bind to FLAG-Rab18. The affinity showed slightly lower compared to PLIN2-FL and PLIN2-251-425 (Fig. 3E). These data demonstrated that Rab18 binds to PLIN2 by its C terminal domain, which is also the LD targeting domain of PLIN2. According to the co-IP results, the PLIN2-251-425 contained the complete structure that interacted with Rab18, and the PLIN2-395-425 may be the specific binding domain for Rab18 with a stabilization structure of PLIN2-370-395. Theses current work reveals that the C terminal domain of PLIN2 is responsible for its LD targeting and the interaction with Rab18, and whether the relationship between the two is competitive or synergistic remains to be studied.

**Figure 3.**
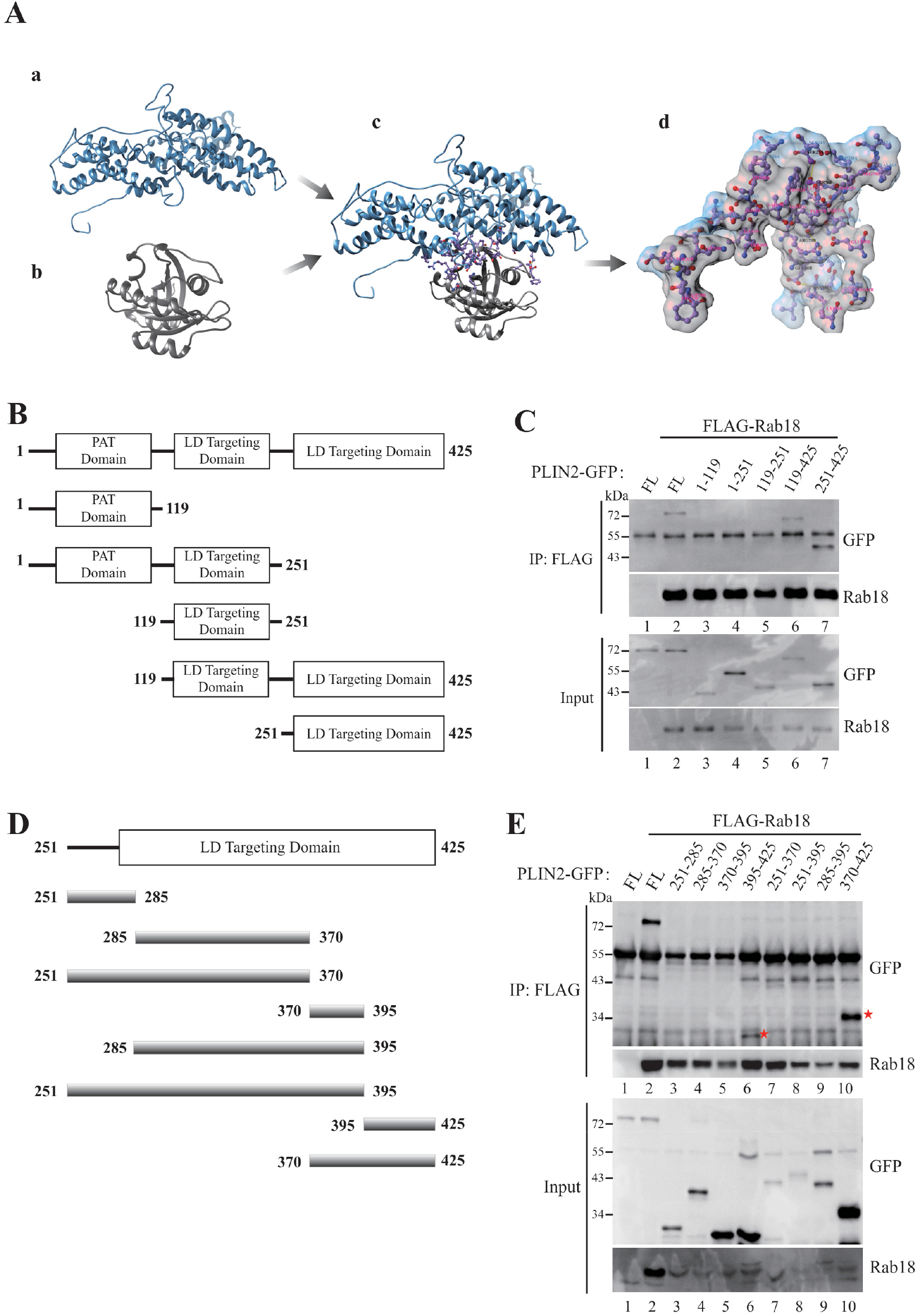
The C terminal domain of PLIN2 is required for its interaction with Rab18. **A**. The PLIN2 and Rab18 protein-protein docking prediction was stimulated by the GrammX model1, which has the highest global energy value. **a**. PLIN2 protein with its most interacting conformation for Rab18. **b**. Rab18 protein with its most probable interacting conformation. **c**. Most probable interaction conformation of PLIN2 and Rab18. d. The most interacting residues of PLIN2 and Rab18. **B**. Line diagram of PLIN2 wild type and its truncations. **C**. HEK293A cells were co-transfected with FLAG-Rab18 and full length (FL) PLIN2-GFP, as well as its truncations. After OA treatment, cell lysates were prepared and performed IP using anti-FLAG M2 beads. **D**. The further schematic representation of PLIN2 C terminus. E. HEK293A cells were cotransfected with FLAG-Rab18 and PLIN2-GFP wild type, as well as its further truncations in panel D, then performed IP using anti-FLAG M2 beads.

**Table 2.** The specific common interacting residues for PLIN2 and Rab18

| Protein | A: PLIN2 | B: Rab18 |
| --- | --- | --- |
| Interacting residues | HIS252, SER249, SER253,<br>GLU385, TYR389, ASP388,<br>ASP384, ASN269, HIS266,<br>GLN270, GLN273, PHE403,<br>PRO402, VAL400, PHE206,<br>HIS256, LEU257, ARG262 | ARG69, GLN67, ARG79,<br>THR108, GLU105, ASN104,<br>ASP100, HIS143, GLU107,<br>LYS142, LEU73, THR72,<br>ARG71, VAL97, ASP94,<br>GLU68, LYS98, ASN101 |

#
#
#

### Rab18 depletion leads to less TAG accumulation and fewer but larger LDs

As a specific LD-translocation of Rab18 occurred during OA induced LD growth and accumulation, we asked about its function on the growing LDs and what role it plays in LD growth and accumulation. To answer these questions, knockdown of Rab18 was conducted by siRNA (Fig. 4B). The TAG accumulation stimulated by OA was significantly suppressed when Rab18 was knockdown (Fig. 4A), which indicated that Rab18 plays an important role in OA induced TAG accumulation.

**Figure 4.**
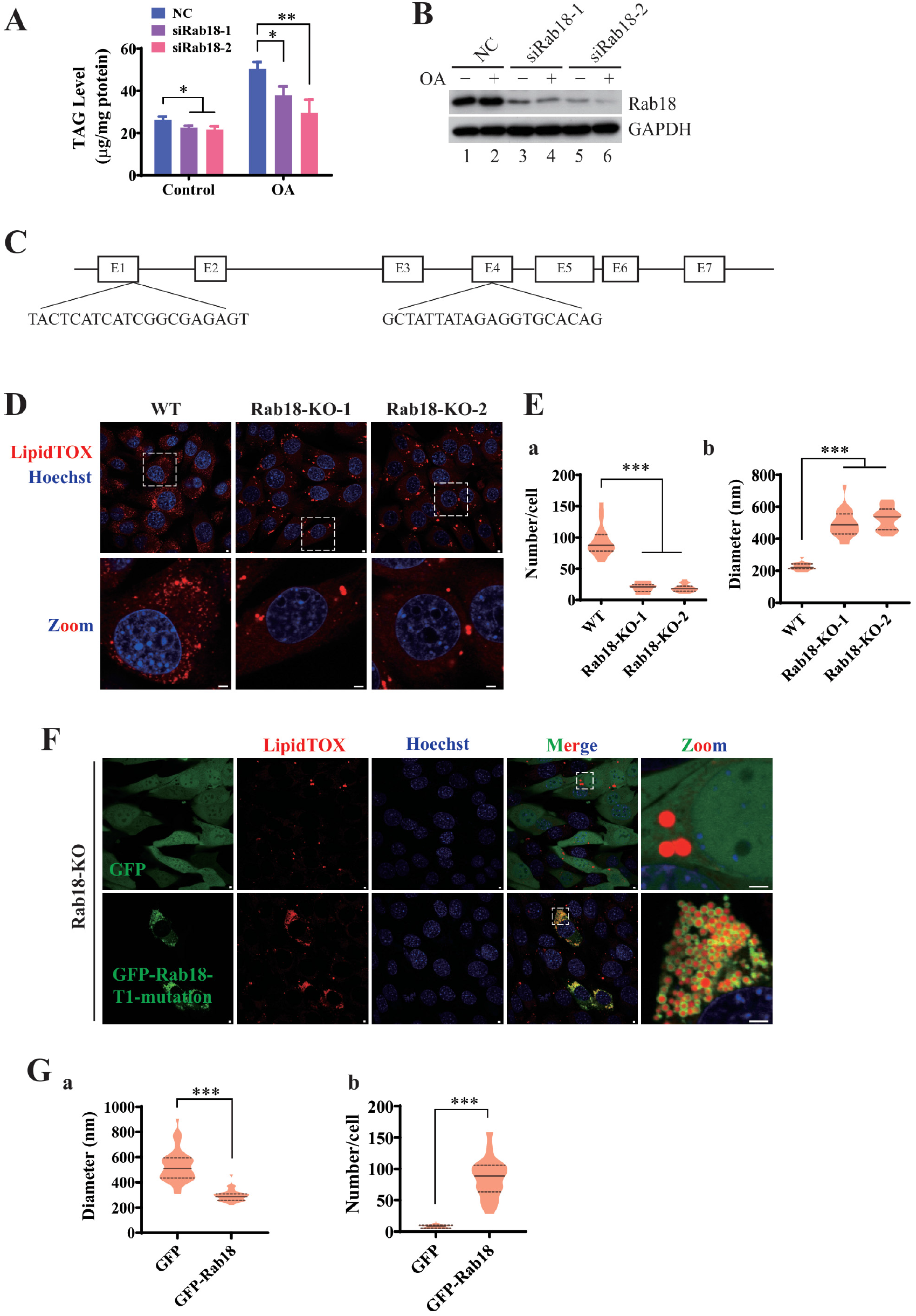
Rab18 depletion leads to less TAG accumulation and less but larger LDs A. C2C12 cells were transfected with two Rab18 siRNA separately, after 48 h, the cells were treated with or without OA for 12 h, and then the whole cell TAG content was analyzed (n=3), and the Rab18 protein expression level was measured by Western bolt (**B**). **C**. CRISPR/Cas9 was used to knock out Rab18 in C2C12 cells. Two target sites for gRNA design were shown. **D**. WT and two Rab18 KO cells were treated with OA for 12 h and LD morphology was detected by confocal imaging. Scale bar=2 μm. **E**. The average size (**a**) and number of LDs per cell (**b**) were analyzed from at least 20 cells each group. **F**. Rab18-KO-1 cells were infected with the virus containing GFP control or GFP-Rab18 T1 mutant, then treated with OA for 12 h. Scale bar=2 μm. **G**. Statistical analysis were calculating from at least 40 cells each group. Data shown as Mean ± SD; *, P≤0.05; **, P≤0.01; ***, P≤0.001.

Then, Rab18 was knocked out in C2C12 cells by CRISPR/Cas9. To ensure the KO efficiency, two pairs of guide RNAs were used to construct two KO cell lines in which Rab18 was mutated in exon 1 and exon 4 respectively (Fig. 4C). Monoclones of Rab18 deletion cells were selected and verified by western blotting (Fig. S1). Consistent with the knockdown cells, LDs in Rab18 null cells (Fig. S1A) showed aberrant biogenesis with OA treatment, which mainly manifested as much less LDs, with a larger size (Fig. 4D). Quantitative analysis of LD size distribution showed that Rab18 deletion resulted in a 2-fold increase of average diameter compared to control cells (Fig. 4E, a). However, the numbers in Rab18 deleted cells were much less than that of control cells with the average number of 20 ± 7 and 19 ± 6 per cell, compared with that of 95 ± 25 per cell (Fig. 4E, b). Furtherly, we reintroduced a knockout-resistant Rab18 mutant into Rab18-KO cells. As a control, overexpression (OE) of GFP did not change the LD phenotype in Rab18-KO cells, however, replenishment of GFP-Rab18 into Rab18-KO cells increased the number of LDs and decreased the size of LDs (Fig. 4F). Quantitative analysis confirmed that reintroduction of GFP-Rab18 could significantly promote more LD formation and at the same time uniform the LD size distributions (Fig. 4G). These results demonstrated that Rab18 plays an essential role in regulating TAG accumulation and LD morphology.

### Rab18-induced TAG accumulation is dependent on the function of ACSL3

As a key factor in OA-stimulated LD formation, Rab18 was translocated from ER to LDs through binding with PLIN2. To further address the regulation mechanisms of the LD associated Rab18 in TAG accumulation and LD dynamic, we constructed FLAG-Rab18 overexpressed stable cell line (Rab18-OE), as well as its constitutively active form (Rab18-Q67L-OE) and dominant negative form (Rab18-S22N-OE) overexpressed stable cell lines. Monoclonal stable cell lines were selected and verified by Western blotting (Fig. 5A). We observed that more TAG was accumulated in Rab18-OE and Rab18-Q67L-OE cells compared with WT cells with OA treatment, which suggested Rab18 may regulate the neutral lipid storage capacity. However, the overexpression of Rab18-S22N had no effect on TAG level, which indicated that GTP binding ability of Rab18 is indispensable for its role in TAG accumulation (Fig. 5B). In support of the elevated TAG accumulation by functional Rab18, the LD phenotype was also analyzed in Rab18-OE cell lines under OA treatment. Strikingly, upregulated expression of Rab18 or Rab18-Q67L was associated with a dramatic increase in LD numbers compared to WT cells (Fig. 5C). It implied that TAG accumulation was associated with increased formation of LDs when Rab18 GTPase activated. In contrast, less but larger LDs were observed in Rab18-S22N-OE cells, which showed similar LD phenotype with Rab18-KO cells (Fig. 5C). It indicated that Rab18 mediated LD formation depends on its GTPase activity. Moreover, the slightly reduced size of LDs suggested the upregulated Rab18 promotes LDs dynamic by increase the number of LDs and the specific surface area. Taken together, Rab18 could increase the TAG accumulation and LDs number, which was regulated by its guanine nucleotide status.

**Figure 5.**
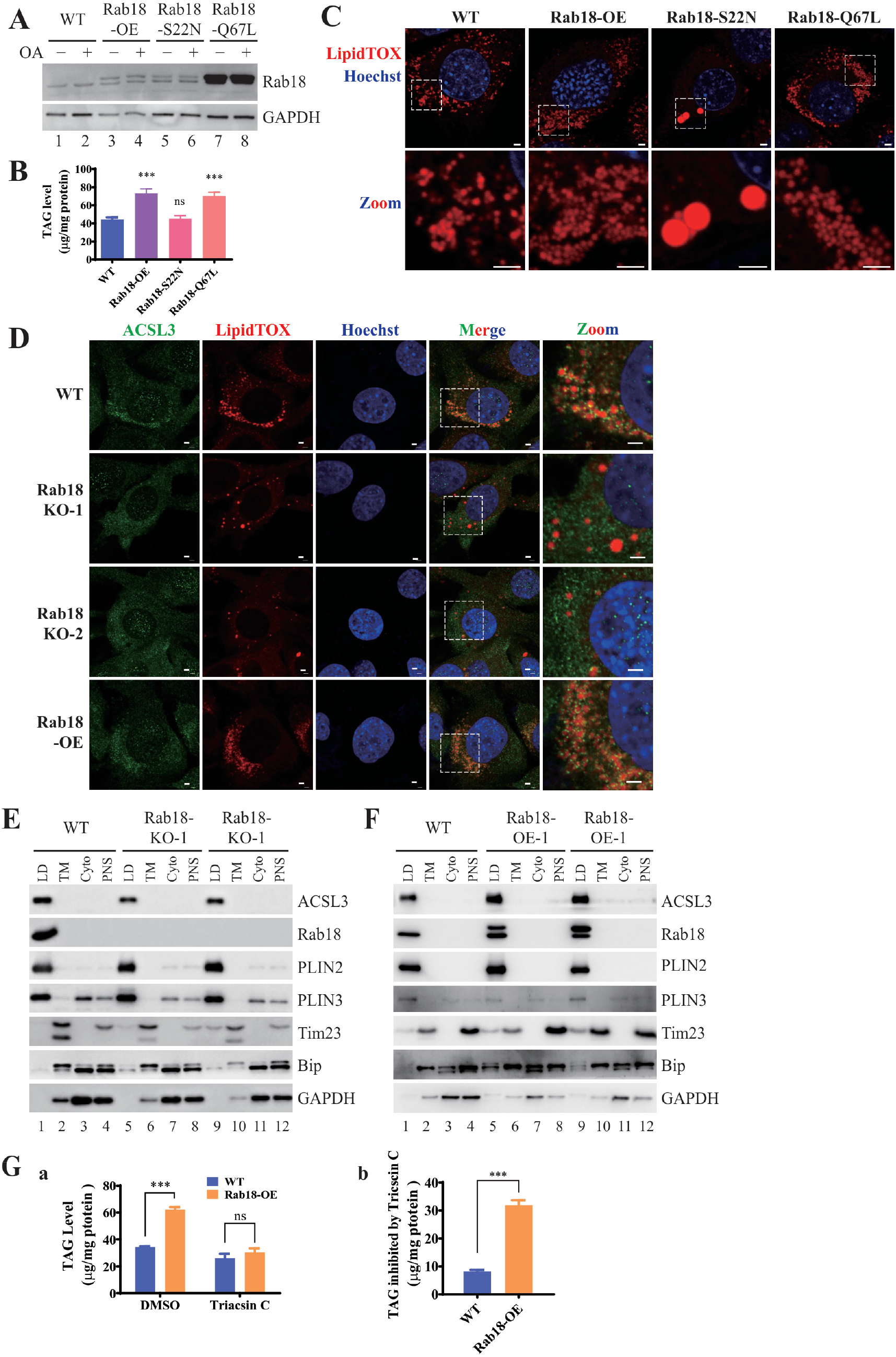
Rab18 induced TAG accumulation is dependent on the LD translocation of ACSL3. **A**. WT, FLAG-Rab18-OE, FLAG-Rab18-S22N-OE and FLAG-Rab18-Q67L-OE C2C12 cells were treated with or without OA, and the Rab18 expression was analyzed by Western blot. **B**. WT, FLAG-Rab18-OE, FLAG-Rab18-S22N-OE and FLAG-Rab18-Q67L-OE C2C12 cells were treated with OA. The TAG content was analyzed and normalized by protein content. n=6. **C**. WT, FLAG-Rab18-OE, FLAG-Rab18-S22N-OE and FLAG-Rab18-Q67L-OE C2C12 cells were treated with OA. The LD morphology was observed by confocal imaging. Scale bar=2 μm. **D**. ACSL3 localization was analyzed in OA-treated WT, two Rab18 KO monoclonal cells and Rab18 OE cells by immunofluorescence with anti-ACSL3. Scale bar=2 μm. **E-F**. LDs were isolated from WT and two Rab18 KO clones (**E**) or two Rab18 OE clones (**F**) after treated with OA. The presence of ACSL3 in LD and other fractions was detected by Western blot. **G**. WT and Rab18 OE cells were treated with OA and Triacsin C for 12 h, then lysed for TAG measurement (**a**). Inhibited TAG production by Triacsin C in WT and Rab18 OE cells was calculated (**b**). Data shown as Mean ± SD, ***, P≤0.001; ns, no significance.

To gain an insight into the mechanism how Rab18 regulate LD dynamic and TAG accumulation, we focus on another Rab18 interacting protein candidate, ACSL3. As listed in table 1, ACSL3 was the most abundant protein interacting with Rab18 in band 1, which was verified by Western blot in Fig. 2B. We next determined the interaction between Rab18 and ACSL3. ACSL3 targets to initial LDs and regulates LD biogenesis[50]. To figure out the effect of Rab18 on the function of ACSL3, we detected the LD localization of ACSL3 in Rab18 KO and OE cells. First, the localization of ACSL3 by immunofluorescence with anti-ACSL3 was imaged in Rab18 KO and OE cells (Fig. 5D). ACSL3 showed punctate spots around LDs in WT cells. However, most green spots were distributed in other membranes other than LD when Rab18 was deleted. Besides, overexpression of Rab18 increased the LD-associated ACSL3 spots. We further purified the LDs from these cells with OA treatment, and found that the content of ACSL3 in LD fraction was decreased in Rab18 KO cells (Fig. 5E and Fig. S1C) and increased in Rab18 OE cells (Fig. 5F and Fig. S1D), however PLIN2 expression was not affected by Rab18 KO. These results indicated that Rab18 induce ACSL3 to localize to LDs. To figure out whether ACSL3 is involved in the promotion of TAG accumulation by Rab18, Triacsin C was used to inhibit the activity of ACSL3. When C2C12 cells were treated with OA and Triacsin C simultaneously, TAG accumulation was significantly blocked in both WT and Rab18 OE cells (Fig. 5G, a). In particular, it was more thoroughly inhibited in Rab18 OE cells (Fig. 5G, b). These results suggested that the LD associated ACSL3 may contribute to the increased TAG accumulation induced by Rab18. In another word, OA-induced Rab18 translocation facilitated TAG and LD accumulation by recruiting ACSL3 to LDs.

**Table 1.** Most abundant proteins analyzed by mass spectrometry in the three bands from FLAG-Rab18 precipitation (in Fig. 2A)

| Sample ID | Proteins | Pep. No. | MW (kDa) |
| --- | --- | --- | --- |
| 1 | Long-chain-fatty-acid--CoA ligase 3 isoform a | 24 | 80.5 |
|  | Long-chain-fatty-acid--CoA ligase 4 isoform 2 | 13 | 74.3 |
|  | UPF0554 protein C2orf43 homolog isoform 1 | 12 | 37.4 |
|  | Sterol-4-alpha-carboxylate 3-dehydrogenase, decarboxylating | 8 | 40.7 |
|  | Perilipin-2 | 8 | 46.6 |
|  | Lanosterol synthase | 8 | 83.1 |
|  | Dehydrogenase/reductase SDR family member 1 | 7 | 34.0 |
|  | Ras-related protein Rab-18 | 7 | 23.0 |
| 2 | Perilipin-2 | 17 | 46.6 |
|  | Probable saccharopine dehydrogenase | 13 | 47.1 |
|  | Ras-related protein Rab-18 | 10 | 23.0 |
|  | Eukaryotic initiation factor 4A-II isoform a | 3 | 46.4 |
|  | Ras-related protein Rab-2A | 3 | 23.5 |
|  | Ras-related protein Rab-3 | 3 | 21.5 |
| 3 | Perilipin-2 | 16 | 46.6 |
|  | UPF0554 protein C2orf43 homolog isoform 1 | 12 | 37.4 |
|  | Sterol-4-alpha-carboxylate 3-dehydrogenase, decarboxylating | 9 | 40.7 |
|  | Dehydrogenase/reductase SDR family member 1 | 6 | 34.0 |
|  | Ras-related protein Rab-18 | 6 | 23.0 |
|  | Glycerol-3-phosphate acyltransferase 6 precursor | 4 | 52.2 |

#

## Discussion

Previous studies have demonstrated that Rab18 is localized to different membrane structures in different cell lines. The LD targeting of Rab18 is identified by several LD proteomics [19, 51, 52], and verified by subcellular fractionation, fluorescence labeling and immune-electron microscopy in hepatocyte, adipocyte and kidney cells[26, 33]. We also detected that Rab18 was recruited from ER to LDs during LD biogenesis and growth in mouse myoblast C2C12 cells (Fig. 1). Mechanisms of protein localization to LDs were classified into several pathways [53]. Rab18 contains a mono-cysteine prenylation motif in its C-terminus which is required for its membrane targeting, but the lipid anchor motif is insufficient to target Rab18 specifically to LDs [27]. For Rab proteins, the correct membrane targeting is achieved by a complex interplay of several factors including lipid anchor motifs, GEFs, effectors and protein-protein interaction [54]. Rab3GAP1/2 is initially reported as a specific Rab18 GEF, which regulates the ER targeting of Rab18 [32]. However, a recent study states that Rab3GAP1/2 also accounts for the LD targeting of Rab18 [38]. Besides, TRAPP II has been previously found to be a GEF for Rab18 and recruit Rab18 onto LDs [39]. Except for direct interaction with lipids of LDs, proteins can target to LDs by interacting with other LD proteins. Based on this point, we focus on LD-associated proteins interacting with Rab18. Two abundant proteins interacted with Rab18 were identified by mass spectrometry analysis, ACSL3 and PLIN2 (Fig. 2 A, Table 1). ACSL3 has a dual localization in ER or on LDs, and it translocated from the ER to LDs during initial LD formation [50]. PLIN2 belongs to perilipin family proteins and is a resident protein of LDs. We further found that PLIN2 depletion significantly reduced the LD localization of Rab18 (Fig.2 D-H). It seems that the LD targeting of Rab18 is enhanced by interacting with PLIN2. Furthermore, the interaction between PLIN2 and Rab18 dependents on the C-terminus domain of PLIN2, which is important for its highest affinity lipid binding [49]. In other words, the C-terminus domain of PLIN2 is not only a LD binding domain, but also a specific Rab18 binding domain. We further found that changes in Rab18 expression level had no effect on the LD localization of PLIN2 in Rab18 KO or OE C2C12 cells (Fig. 5E and F), which suggests that the relationship between LD targeting and Rab18 binding of PLIN2 is not mutually exclusive. However, there is another study reported that PLIN2 and Rab18 are localized to different population of LDs and Rab18 overexpression reduces the amount of PLIN2 in HepG2 cells [26]. The disparity may attribute to different cell types. The LD phenotype is also different between the cell lines. It is said that overexpression of Rab18 does not increase the number or the total volume of LDs in HepG2 [26]. Xu et al. [38] also observe a specific role of Rab18 in controlling lipid storage and LD growth in 3T3L1 and TM-2 cells, but not in AML12, HeLa, Cos7 and HEK293T cells. In addition, Jayson et al. has demonstrated that Rab18 is not a general and necessary component of the protein machinery involved in LD biogenesis or turnover in a human mammary carcinoma cell line[55]. Nevertheless, our results showed that overexpression of Rab18 and its GTP form, Rab18-Q67L significantly increased the number of LDs in C2C12 myoblasts (Fig. 5C), LDs formation was totally blocked when Rab18 was deleted or mutated to the dominant-negative form (Fig. 4D, 5C).

Aberrantly large LDs in Rab18 deficient cells is consistent in multiple studies. We also detected larger but less LDs in Rab18 KO C2C12 cells, which can be recovered by replenishment of Rab18 (Fig. 4F). Moreover, TAG accumulation was reduced in Rab18 KO cells. It implies abnormal LD formation and/or LD fusion caused by loss of Rab18. A recent study proposes that Rab18 deficiency don’t affect nascent LD formation and LD fusion, but controls the growth of nascent LDs into mature LDs[38]. However, it is still difficult to elucidate how the larger LDs form in the Rab18 depleted cells. On the other hand, Rab18 OE cells accumulated much more normal size LDs (Fig. 5C). The TAG content accumulating while the LDs “still keep fit”, which suggests that “high metabolic rate” may be a good explanation for Rab18 overexpressed cells, with higher specific surface area. Manipulation of functional Rab18 expression could change TAG accumulation and determine LD number and size. However, it is still a question whether the increase of TAG content results from more TAG synthesis or less TAG hydrolysis.

Rab18 may regulate lipid homeostasis by promoting the interaction between LD and ER [26, 38]. We identify that another protein, ACSL3 may also involve in this process. ACSLs are key enzymes that provide acyl-CoA for lipid metabolism. ACSL3 can localize to both ER and LD and promote LD biogenesis and fatty acid uptake [50, 56]. The interaction between Rab18 and ACSL3 may promote the LD association of ACSL3, since recruitment of ACSL3 onto LDs was defective in Rab18 depleted cells and enhanced in Rab18 OE cells (Fig. 5D and F). Similarly, it has been studied that Rab18 activity is also required for the association of ApoB to LDs [57]. Moreover, when the activity of ACSL3 was inhibited by Triacsin C, the increased TAG in Rab18 OE cells was completely blocked (Fig. 5G). These data raise the possibility that LD-associated Rab18 reinforces the LD targeting of ACSL3, which facilitating fatty acids and TAG transforming.

In conclusion, we discovered a LD protein complex PLIN2/Rab18/ACSL3 regulating LD homeostasis in myoblasts. PLIN2 facilities and stabilizes the LD targeting of Rab18, which in turn promotes ACSL3 recruitment onto LDs for acyl-CoA accumulation. The TAG amount and LD size growth were regulated by this complex, which mostly depends on the activity of Rab18.

## Conflict of Interest

All authors declared no competing interests.

## Author Contributions

P. L. conceived the project. Y. D., C. Z., S. X. and T. B. conducted the experiments. P. L., Y. D., designed the experiment and analyzed the data. H. M. performed the bioinformatics data. P. L, C. Z., S. Z., and Y. D. wrote the manuscript.

## Acknowledgements

The authors thank Dr. Yan Teng for the assistance of confocal microscopy experiments, Dr. Zhensheng Xie for the mass spectrometry analysis.

This work was supported by the National Key R&D Program of China (Grant No. 2016YFA0500100, 2018YFA0800700 and 2018YFA0800900), National Natural Science Foundation of China (Grant No. 91857201, 91954108, 31671402, 31671233, 31701018 and U1702288). This work was also supported by the “Personalized Medicines——Molecular Signature-based Drug Discovery and Development”, Strategic Priority Research Program of the Chinese Academy of Sciences, Grant No. XDA12040218.

## Abbreviations

ER: Endoplasmic reticulum
IP: Immunoprecipitation
KO: Knock out
LD: Lipid droplet
OA: Oleic acid
OE: Overexpression
WT: Wild type

## Notes

### Competing Interest Statement

The authors have declared no competing interest.

