## Supplementary material for "Rab18 Binds PLIN2 and ACSL3 to Mediate Lipid Droplet Dynamics": Figure S1

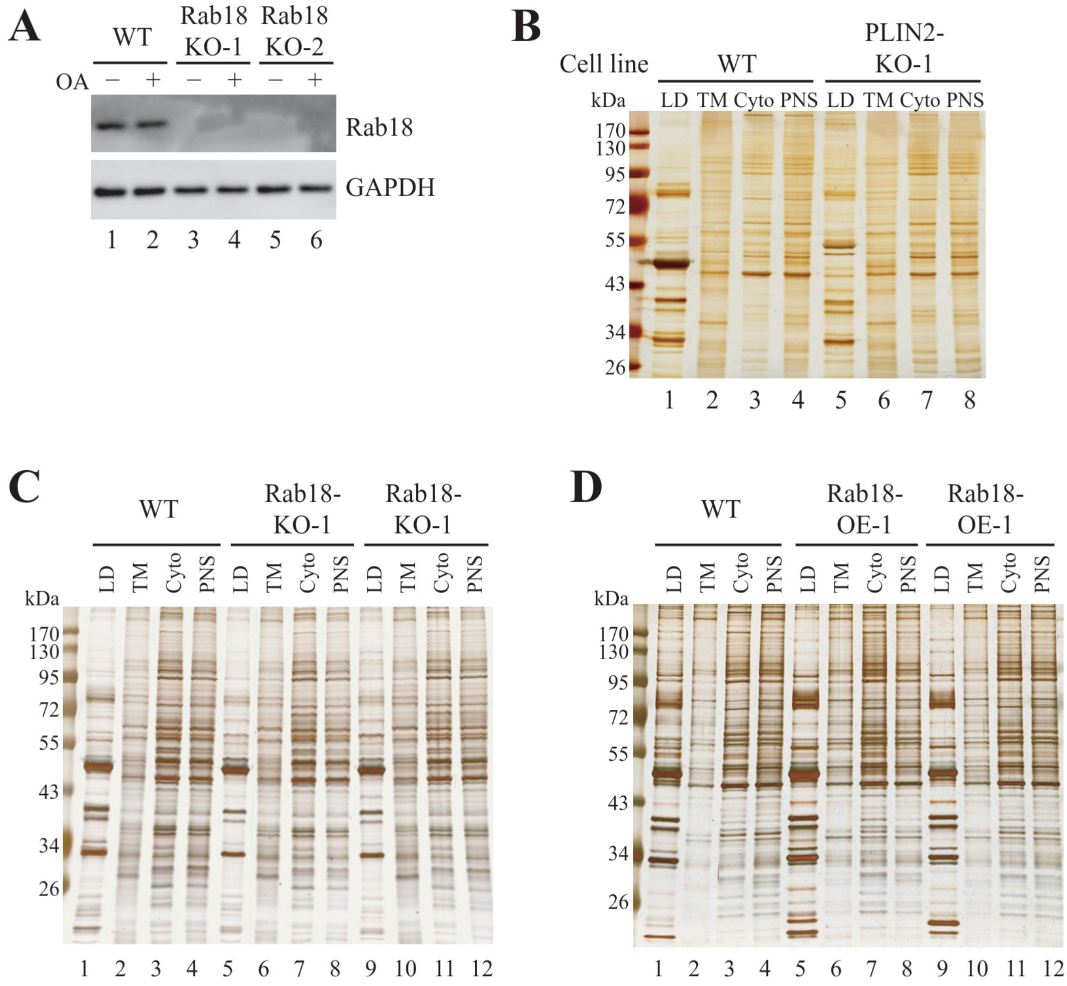


**Figure S1. Rab18 and PLIN2 KO protein expression control**

**A**. The Rab18 protein expression level in the WT and Rab18 KO monoclones with or without OA treatment was measured by Western bolt. **B**. Protein profile of different cell fractions by silver staining in WT and PLIN2-KO cells, as in Fig.2E. **C**. Protein profile of different cell fractions by silver staining in WT and Rab18-KO cells, as in Fig.5E. **D**. Protein profile of different cell fractions by silver staining in WT and Rab18-OE cells, as in Fig.5F.
